# Antiviral drug synergy and mutational signatures in different epithelial cell models of RSV and hPIV infection

**DOI:** 10.64898/2026.01.13.699296

**Authors:** Samuel Ellis, Amy I. Jacobs, Shengyuan Zhang, Laura Buggiotti, Maximillian Woodall, Jila Ajeian, Helena Tutill, Charles Miller, Macheala Palor, Angelika Kopec, Elizabeth K. Haughey, Hongxia Ma, Mia Tomlinson, Rachel Williams, Chris O’Callaghan, Joseph F. Standing, Claire M. Smith, Judith Breuer

## Abstract

Despite the huge global health burden presented by respiratory viruses, effective broad-spectrum antiviral therapeutic options remain limited. Here we evaluated the antiviral activity of four RNA-dependent RNA polymerase (RdRp) inhibitors, remdesivir, ribavirin, favipiravir, and molnupiravir, against respiratory syncytial virus (RSV) and human parainfluenza (hPIV) using monotherapy or dual-drug combinations with epithelial cell lines and primary human airway culture models.

Remdesivir showed the greatest potency across both viruses, while ribavirin and favipiravir also demonstrated inhibition. Molnupiravir was active against RSVA but not hPIV3. Several dual-drug combinations, including remdesivir–favipiravir, remdesivir–molnupiravir and favipiravir–molnupiravir, produced marked synergy against RSVA, and more limited synergy for hPIV3. Antiviral efficacy was validated in primary airway epithelial cultures, where effective concentrations preserved epithelial integrity and attenuated viral disruption of ciliary function. Across both viruses, increasing antiviral exposure was associated with dose-dependent signature mutagenesis. Importantly, antivirals induced significantly higher RSVA mutation burden in the primary airway model.

These findings highlight the therapeutic potential of RdRp inhibitor combinations for RSVA and hPIV3, provide mechanistic insight through antiviral-related mutational signatures and demonstrate advantages of the primary human airway culture model for development of effective multi-drug regimens and broad-spectrum antiviral preparedness.

**Graphical abstract:** 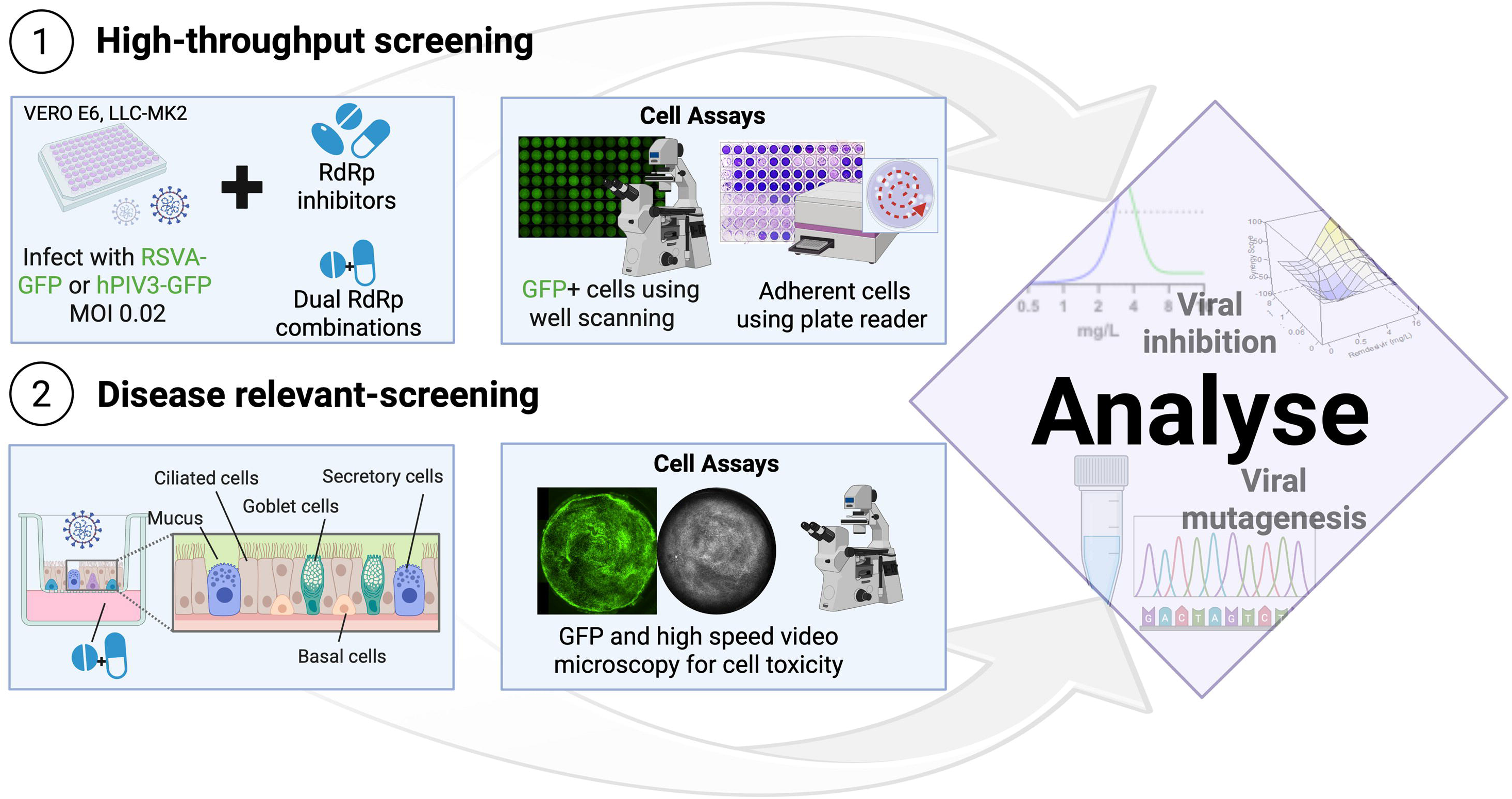

## Introduction

Respiratory viral infections are a major cause of global morbidity and mortality. Respiratory syncytial virus (RSV) and human parainfluenza (hPIV) are leading causes of lower respiratory tract infection and viral pneumonia^1-3^, particularly in infants, the elderly and immunocompromised patients^4,5^. RSV alone infects approximately 33.8 million children below the age of five annually^6^, while PIV contributes substantially to paediatric hospital admissions. More broadly, respiratory RNA-viruses have driven major outbreaks and pandemics, underscoring the need for effective antiviral strategies.

Current post-exposure treatments rely on symptom management, and there remains an unmet need for quick and effective therapies, especially in severe cases such as respiratory viral infections in immunocompromised patients. One promising approach is targeting the RNA-dependent RNA polymerase (RdRp) protein. The RdRp is a highly conserved enzyme across RNA virus species that is essential for viral replication^7^. Interest in RdRp inhibitors increased during the COVID-19 pandemic with efforts to repurpose existing nucleoside analogues^8,9^. Ribavirin, originally developed for hepatitis C virus, is licensed for RSV and shows *in vitro* activity against hPIV, but clinical efficacy is inconsistent and limited by toxicity, cost and safety concerns^10-17^. Other RdRp inhibitors such as remdesivir, favipiravir and molnupiravir were initially developed for influenza and severe acute respiratory syndrome (SARS)^7,18^. All three have subsequently been investigated for potential efficacy against other RNA viruses, most notably being repurposed for SARS-CoV-2 during the COVID-19 pandemic ^18-20^.

Alternatively, drug combination may enhance antiviral efficacy and tolerability through additive or ideally synergistic inhibitory effects^21^ and has been highly successful in other viral diseases, most notably HIV^22^. Within the field of respiratory RNA viruses, combination therapy has been suggested as an important approach for improving influenza treatments, particularly for mitigating the emergence of resistance^23^. In recent years, the use of multiple antivirals has been investigated for SARS-CoV-2, with many combinations demonstrating synergy *in vitro* and *in vivo*^9,24,25^. Broad-spectrum combinations with differing viral susceptibilities may be particularly advantageous for vulnerable populations such as immunocompromised patients^26^. Robust evaluation of such strategies, however, requires experimental systems that reliably capture both antiviral activity and host epithelial responses across multiple respiratory viruses.

Most antiviral studies rely on immortalised epithelial cell lines as an infection model, which offer practical advantages such as ease of culture, reproducibility and suitability for higher-throughput screening. However, these models incompletely represent the complexity of the human airway epithelium, and different respiratory viruses frequently require distinct cell lines to achieve efficient replication or quantifiable cytopathology, limiting standardisation and complicating direct comparisons of antiviral efficacy across viral species^27^.

Primary human airway culture models provide a more physiologically relevant approach. When differentiated at an air–liquid interface, these cultures recapitulate the pseudostratified architecture of the human airway, incorporating ciliated and secretory cell populations, functional tight junctions and mucus secretion. Importantly, this model supports infection by multiple respiratory viruses within a unified experimental model^28^, enabling direct evaluation of antiviral efficacy, epithelial toxicity and virus-induced functional disruption, such as impaired ciliary beating.

Here, we address these gaps by systematically evaluating the *in vitro* efficacy of selected RdRp inhibitors, both as monotherapies and in combination, against RSV and hPIV. Using conventional epithelial cell lines alongside primary human airway epithelial cultures, we aim to define antiviral potency, synergistic interactions and epithelial outcomes in models that better reflect human respiratory infection, thereby informing the development of effective and broadly applicable antiviral strategies.

## Materials and Methods

### Viral strains and culture

Two laboratory strains of GFP-tagged RSV A2 were used. RSV-GFP1 was purchased from ViraTree (product code R121) and was titrated and used directly for all RSVA antiviral assay infections unless otherwise specified. Some supplementary and optimisation assays in this study also used a second GFP-RSV strain, propagated in-house from viral stocks originally generated and provided by collaborators^29^. All hPIV experiments used a GFP-tagged hPIV3 strain purchased from ViraTree (product code P323). All the above viral stocks were aliquoted, snap frozen and stored at -70°C or -150 °C until experimental use.

For RSVA propagation, T175 flasks (Corning) of HEp-2 cells were infected at a multiplicity of infection (MOI) of 0.01 in serum-free VP-SFM media (Thermo Fisher) supplemented with 4 mM L-glutamine and 0.5% v/v Pen/Strep until approximately 80% of cells demonstrated cytopathic effects (CPE) and/or GFP fluorescence (approximately 5 days post inoculation). Cells were harvested and lysed by sonication and freeze thaw cycle at -80°C. After centrifugation (1600 x g for 10 minutes), virus-containing supernatant was purified and concentrated using 100 kDa Vivaspin-20 ultrafiltration tubes (2500 x g, 2-4 hours).

RSVA viral stock titration was performed by TCID50 assay and syncytial plaque forming unit (SPFU) counting by GFP fluorescence, both using serial dilution inoculation of VeroE6 cells on 96-well culture plates. TCID50 CPE functional titre was calculated by the Spearman & Kärber algorithm as described^30^. TCID50 plates were further imaged on a Nikon Ti-E microscope, using GFP-signal to count syncytial plaques per well to calculate SPFU/ml titre. The latter titre was used for all MOI calculations for subsequent experiments in this study, unless otherwise stated.

### Epithelial Cell Lines

African green monkey kidney cell line Vero E6 (ATCC: C1008-CRL-1586) was provided and authenticated by The Francis Crick Institute, London, UK, for use in this study. Vero E6 cells were routinely maintained in Dulbecco’s Modified Eagle Medium (DMEM) supplemented with 10% fetal calf serum (FCS, Thermo Fisher) and 1× penicillin/streptomycin (Sigma-Aldrich). Human lung adenocarcinoma cell line Calu-3 (HTB-55, ATCC) and rhesus monkey kidney cell line LLC-MK2 (CCL-7, Caltag-Medsystems Ltd) were maintained in DMEM supplemented with 10% fetal bovine serum (FBS, Thermo Fisher) and 1× penicillin/streptomycin. All cells were incubated at 37°C and 5% CO2 in a humidified incubator, with weekly passaging and media replacement 1-3 times a week as needed.

### Primary airway epithelial culture

The air-liquid interface (ALI) culture model for human primary airway epithelium was performed as previously described, from a nasal epithelial brush biopsy collected from a healthy paediatric donor (3-year-old female) under ethical approval for the study provided by the UCL Living Airway Biobank (REC reference 14/NW/0128)^31^.

In brief, basal epithelial cells were first expanded using co-culture with mitomycin-inactivated 3T3-J2 fibroblasts, then separated from feeder cells using differential trypsinization. Epithelial cells were seeded onto collagen I-coated, 24-well plate semi-permeable membrane supports (Transwell, 0.4 µm pore size, Corning). To induce differentiation, apical fluid was removed to expose cells to air (5% CO2), and basolateral media was replaced with PneumaCult ALI differentiation media (STEMCELL Technologies), exchanged three times weekly. These ALI cultures were maintained for a minimum of 4 weeks for differentiation before downstream experiments. Transepithelial electrical resistance (TEER) was regularly measured using an EVOM2 resistance meter (World Precision Instruments) to indicate differentiation and monitor epithelial barrier function.

### Antiviral Drugs

The following antiviral drug compounds were acquired for this study: remdesivir (Bio-Techne), favipiravir (Fisher Scientific), EIDD-1931 (active metabolite of molnupiravir; Sigma-Aldrich), ribavirin (Cambridge Bioscience). These were purchased as lyophilised product and resuspended in DMSO at a standard stock concentration of 10 mg/mL. The drug stocks were stored as single-use aliquots at –20°C or 4°C, per manufacturer recommendations.

### Cell line model for assessing antiviral efficacy

Cell line assays in 96-well format were adapted from our previous SARS-CoV-2 antiviral study^9^, with the following modifications for optimisation with these viruses. For RSVA assays, VeroE6 cells were seeded the day before infection at a density of 1-2x10^4^ cells per well in a black walled, clear bottomed 96 well culture plate (Corning Costar). Six plates were seeded per experiment, for triplicate plates measuring antiviral infection inhibition (with virus) and antiviral drug cytotoxicity (without virus). For infection, culture media was removed and viral inoculum added at MOI of 0.02 (or media control for cytotoxicity plates) in 50 µL of serum-free DMEM and incubated at 37°C and 5% CO_2_ for 1-2 hours. Each well was then completed to 200 µL volume containing experimental antiviral drug concentration and final concentration of 5% FCS.

Assay plates were incubated at 37°C and 5% CO_2_ until experimental endpoint (typically 7 days post-inoculation unless otherwise specified). The plates were regularly monitored for CPE by inverted light microscope, and at chosen experimental timepoints all wells were recorded for GFP fluorescence on a Nikon Ti-E microscope using an automated NIS-Elements JOBS (Nikon) module. At experiment endpoint, media was collected for downstream applications including PCR and sequencing, and remaining cells were fixed and stained using 4% paraformaldehyde and 0.1% v/v crystal violet (Sigma). Excess stain was washed off with water and plates air-dried, before crystal violet staining was quantified by absorbance at 595 nm for each well using Spiral Averaging on a FLUOstar Omega plate reader (BMG Labtech).

The same methodology was applied for hPIV3 assays, but with the use of the cell line LLC-MK2 and the infection media containing 5% FBS.

### Statistical analysis of antiviral efficacy

Analysis of the antiviral efficacy was performed as previously described^9^. In brief, the dependent variable for the statistical analysis of antiviral effect was the percentage inhibition read from the optical density at 595 nm from crystal violet staining of remaining cells. Statistical analyses were performed in R (version 4.4.2) including calculation of the variables effective dose for 50 and 90% inhibition (EC50/EC90) for monotherapies, and the SynergyFinder version 3.14.0 package for R was utilised for drug combination analysis. The R code is available on GitHub at https://github.com/ucl-pharmacometrics/RSV-PIV-in-vitro-analysis.

### Primary air-liquid interface (ALI) cell culture model for assessing antiviral efficacy

Fully differentiated primary epithelial cultures were grown at air-liquid interface in 24-well Transwell culture plates as described above. Prior to infection, the apical cell surface was washed with PBS for 10 minutes to remove excess mucus, and any pre-infection TEER measurements or microscopy performed. Viral inoculum (or media only mock control) was added to the apical Transwell at an MOI of 0.05 in 200 μL of PneumaCult media and incubated at 37°C and 5% CO2 for 1 hour. The inoculum was removed, and the basolateral media replaced with 600 μL of PneumaCult media containing chosen antiviral drug concentrations (or media only for controls), in triplicate replicate wells per experimental condition. The cultures were incubated at 37°C and 5% CO2 until experimental endpoint at 6-7 days post-infection (dpi), with TEER measurements and microscopy for GFP fluorescence or cilia beat frequency analysis performed at approximately 2-day intervals. At endpoint, apical washes were collected from all wells by incubating 200 μL PBS for 10 minutes and stored for downstream PCR and sequencing analysis. All cultures were fixed using 4% PFA for 30 minutes and stored submerged in PBS + 0.02% Sodium Azide for downstream immunofluorescent staining and confocal microscopy.

### Cilia beat frequency (CBF) analysis

Cilia beat measurements from the primary airway model were performed at interval and endpoint timepoints using an environmental chamber (5% CO2, 37 °C) connected to an inverted microscope system (Nikon Ti-E). Image acquisition was automated using NIS-Elements JOBS (Nikon) module for fast time-lapse (100 fps) recording. CBF was determined from timelapse files by first extracting the average pixel intensities using ImageJ (NIH) and, secondly by performing a fast Fourier transformation (FFT) on this data using R; 6400 regions of interest (ROI), each with an area of 16.8µm² were analysed per file (pixel resolution = 0.32µm). CBF was computed using ciliR software as described previously^32^.

### qRT-PCR

Culture supernatant samples collected for downstream analysis were stored at -70°C, with dilution at 1:1 in DNA/RNA shield (Zymo Research) for longer-term storage. RNA extraction was performed using the Qiagen Viral RNA mini kit (Qiagen) according to the manufacturer’s instructions. For qRT-PCR, 4 µL of extract was combined with 16 µL of master mix (Takara One Step PrimeScript III and primer-probes for PIV 1/2/3 and RSV A/B) and manually loaded onto a MicroAmp™ EnduraPlate™ Optical 384-Well Plate. The qRT-PCR was performed using a QuantStudio 5 real-time PCR system, and negative template control and positive plasmid controls for RSV A/B and PIV types 1/2/3 were included in each run to control for contamination and amplification, respectively.

### Library preparation and sequencing

Viral whole genome sequencing was carried out by bait capture with biotinylated RNA oligonucleotides used in the Agilent SureSelect^XT^ (SSXT) protocols. A design targeting Parainfluenza 1-3, RSV A and B and Metapneumovirus was designed in-house and synthesised by Agilent Technologies.

11 µl RNA extract was used in first-strand cDNA synthesis with random primers and SuperScript IV (Thermo Fisher Scientific) according to the manufacturer’s instructions. Second-strand cDNA was synthesised using the NEBNext Ultra II Non-Directional RNA Second Strand Synthesis Module (New England BioLabs) according to the manufacturer’s instructions. Double-stranded cDNA was purified using AMPure XP Beads (Beckman Coulter) with a 40 µl elution volume and was quantified using a high-sensitivity dsDNA Qubit kit (Fisher Scientific).

Double stranded cDNA (bulked with male human gDNA (Promega) if required) was sheared using a Covaris E220 focused ultra-sonication system (42 seconds, PIP 75, duty factor 10, cycles per burst 1000). End-repair, non-templated addition of 3’ poly A, adaptor ligation, pre-capture PCR, hybridisation, post-capture PCR and all post-reaction clean-up steps were performed using the SureSelect^XT^ Low Input kit (Agilent Technologies) with minor modifications to the manufacturer’s protocol to account for variable pathogen loads. Quality control steps were performed using the 4200 TapeStation (Agilent Technologies). Samples were sequenced using the NextSeq 500 platform (Illumina). Run data were demultiplexed and converted to fastq files using Illumina’s BCL Convert Software v3.7.5.

### Sequencing analysis

The raw fastq reads were adapted, trimmed and low-quality reads removed using the fastp algorithm. The reads were aligned against the RSVA reference genome (KX765960.1) and the hPIV3 reference genome (NC_001796.2); consensus sequences were then called at a minimum of 20X coverage. The entire processing of raw reads to consensus was carried out using nf-core/viralrecon pipeline^33^. Only samples producing genomes with at least 90% genome coverage at 10X sequencing depth were kept for further analysis. Variant calling was done using the iVAR algorithm using the metagenomic protocol ^34^. Minority variants were filtered if the total coverage at that position was less than 200×, with at least 20 reads in total and at least 5 independent reads with no strand-bias supporting each of the main two alleles, and minimum allele frequency threshold was 5%. Antiviral induced mutations were identified by comparing mutations (nucleotide) from virus only positive control samples with sequences from the antiviral drug challenge conditions. General data processing was carried out in R 4.2.0, figures were made using the ggplot2 package^35^.

### Immunofluorescent staining and confocal microscopy of primary airway model

For immunofluorescence confocal imaging, ALI cultures were fixed using 4% (v/v) paraformaldehyde for 30 min, permeabilized with 0.2% Triton X-100 (Sigma) for 15 min and blocked using 5% goat serum (Sigma) in PBS for 1 hour. The cultures were then incubated overnight at 4 °C with primary antibody dilutions prepared in 5% goat serum in PBS with 0.1% Triton X-100. The primary antibodies used included mouse anti-MUC5AC (Sigma-Aldrich, MAB2011, diluted 1:500), Goat Recombinant Anti-alpha Tubulin (acetyl K40) antibody (Abcam, EPR16772, diluted 1:100), Rabbit Anti-GFP antibody (Abcam, EPR14104, diluted 1:100).

Following primary antibody incubation, cultures were washed and then incubated with secondary antibodies diluted in 1.25% donkey serum in PBS with 0.1% Triton X-100 at room temperature for 1 hour the following day. The secondary antibodies included Donkey Anti-Mouse Alexa Fluor® 594 preadsorbed (Abcam, ab150112, 1:500), Donkey anti-rabbit Alexa Fluor® 488 preadsorbed (Abcam, ab96919, 1:500), Donkey anti-goat rabbit Alexa Fluor® 647 preadsorbed (Abcam, ab15013, 1:500).

After the secondary antibody incubation, cultures were stained with Alexa Fluor 555 phalloidin (ThermoFisher, A34055, 4 μg/ml) for 1 hour and DAPI (Sigma, 2 μg/ml) for 15 mins at room temperature to visualize F-actin and nuclei, respectively. Samples were washed three times with PBS containing 0.1% Tween 20 after each incubation step.

Imaging was carried out using an LSM710 Zeiss confocal microscope, and the resulting images were analysed using Fiji/ImageJ v.2.1.0/153c54.

## Results

### *In vitro* efficacy of remdesivir, favipiravir, molnupiravir or ribavirin as monotherapies against RSVA and hPIV3

We first optimised the infection assays for RSVA and hPIV3 to achieve sufficient infection progression to investigate antiviral efficacy. Infection of both viruses was assessed for three epithelial cell lines (VeroE6, Calu3 and LLC-MK2) using two readouts: endpoint cytopathic effect (CPE) by crystal violet staining of surviving cells, and fluorescence intensity generated by GFP-tagged viral replication.

For RSVA, only VeroE6 cells supported measurable infection by both CPE and GFP signals. For hPIV3, LLC-MK2 cells were permissive to infection, whereas Calu3 and VeroE6 cells demonstrated little to no detectable infection, even at higher MOI. An MOI of 0.02 and a 7-day incubation were selected as optimal for the subsequent antiviral assays, using VeroE6 for RSVA and LLC-MK2 cells for hPIV3 assays **[Figure 1A]**. The results for other timepoints and MOIs for all cell line and virus combinations are summarised in **Supplementary Figure 1**.

**Figure 1.**
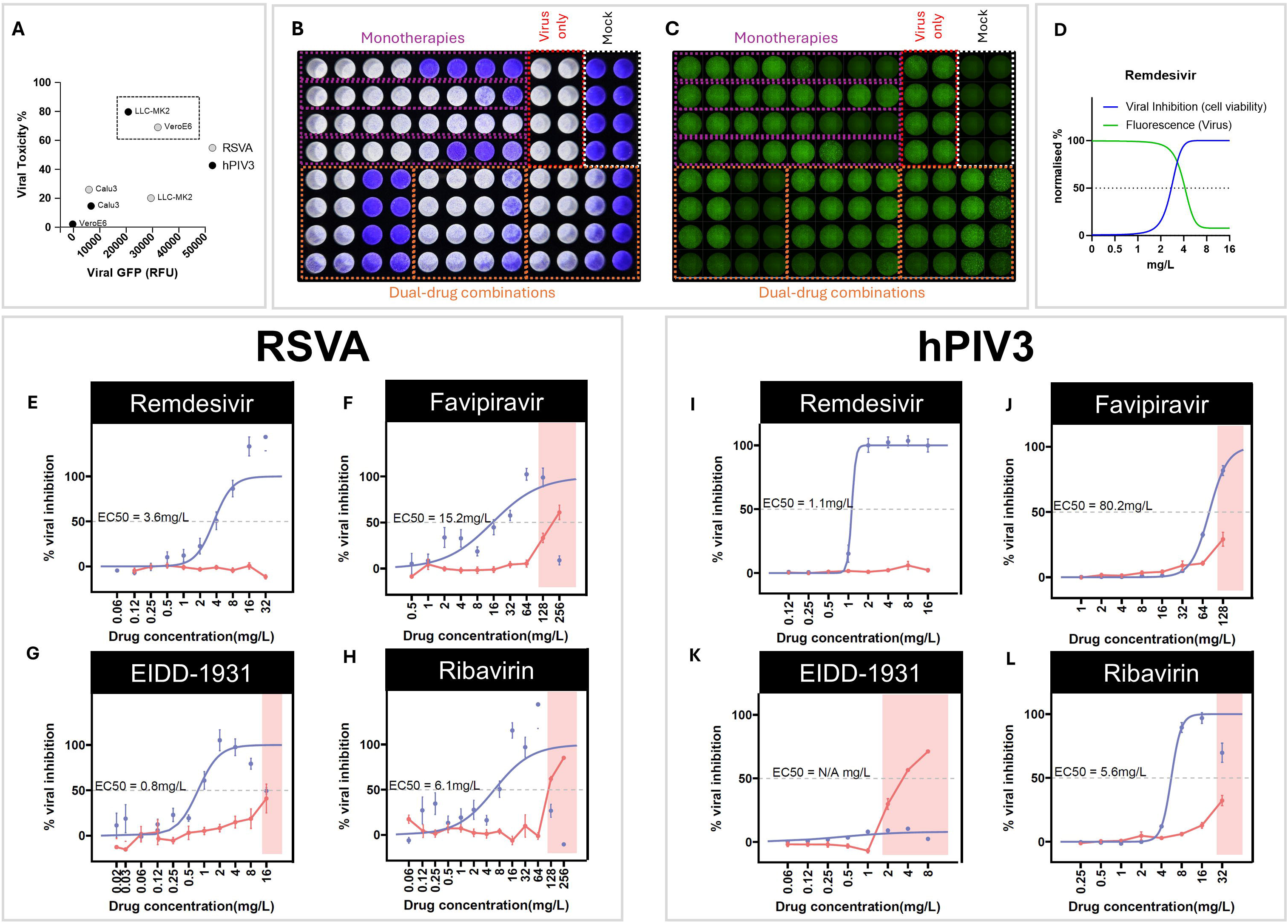
Assay schematic and monotherapy antiviral efficacy. A. Comparison of cell toxicity versus viral-GFP intensity (relative fluorescence units) at 7 days post-infection with RSVA or hPIV3 (both at MOI 0.02) with the epithelial cell lines VeroE6, Calu-3 or LLC-MK2 (n=3). Dashed box indicates cell lines chosen for subsequent antiviral assays. B-C. Representative images of the antiviral infection assay in 96-well format demonstrating dynamic range, here showing RSV infection of VeroE6 cells with and without single or dual-drug antiviral treatments. Cytopathology is visualised by endpoint crystal violet staining of adherent cells (B), whilst the fluorescence micrograph of same representative assay plate (C) demonstrates visualisation of GFP-tagged RSVA infection. D. Relationship between cell viability (quantified by crystal violet absorbance) and viral GFP fluorescence, as normalised against virus-only and mock controls. Representative dose response curves are modelled on data for remdesivir against RSVA. E-L. Dose response curves for monotherapy concentrations of remdesivir (E&I), favipiravir (F&J), EIDD-1931 (G&K), and ribavirin (H&L) tested against RSVA (E-H) or hPIV3 (I-L) infection. The viral inhibition % model is shown as the blue line, while cytotoxicity (as a % reduction of adherent cells) is indicated by the red line. The shaded box indicates drug concentrations with cytotoxicity exceeding 20%. (n=5 for RSVA, n=3-4 for hPIV3)

To determine the monotherapy efficacy of these antivirals, infection assays were performed using serially diluted drug concentrations selected to bracket clinically achievable values: remdesivir (0.06-32 mg/L; 0.1-53.1 μM), favipiravir (0.5-256 mg/L; 3.2-1629 μM), EIDD-1931 (0.02-16 mg/L; 0.08-61.7 μM), and ribavirin (0.06-256 mg/L; 0.2-1048 μM). Endpoint CPE quantification at day 7 **[Figure 1B]** was used to calculate relative viral inhibition versus virus-only and mock controls, from which data a dose-response curve was modelled to determine EC50/EC90 values for each drug-virus interaction **[Table 1]**. Wells were also scanned for GFP fluorescence intensity, with drug-induced viral inhibition reflected in reduced fluorescence intensity **[Figure 1C&D]**.

**Table 1.**
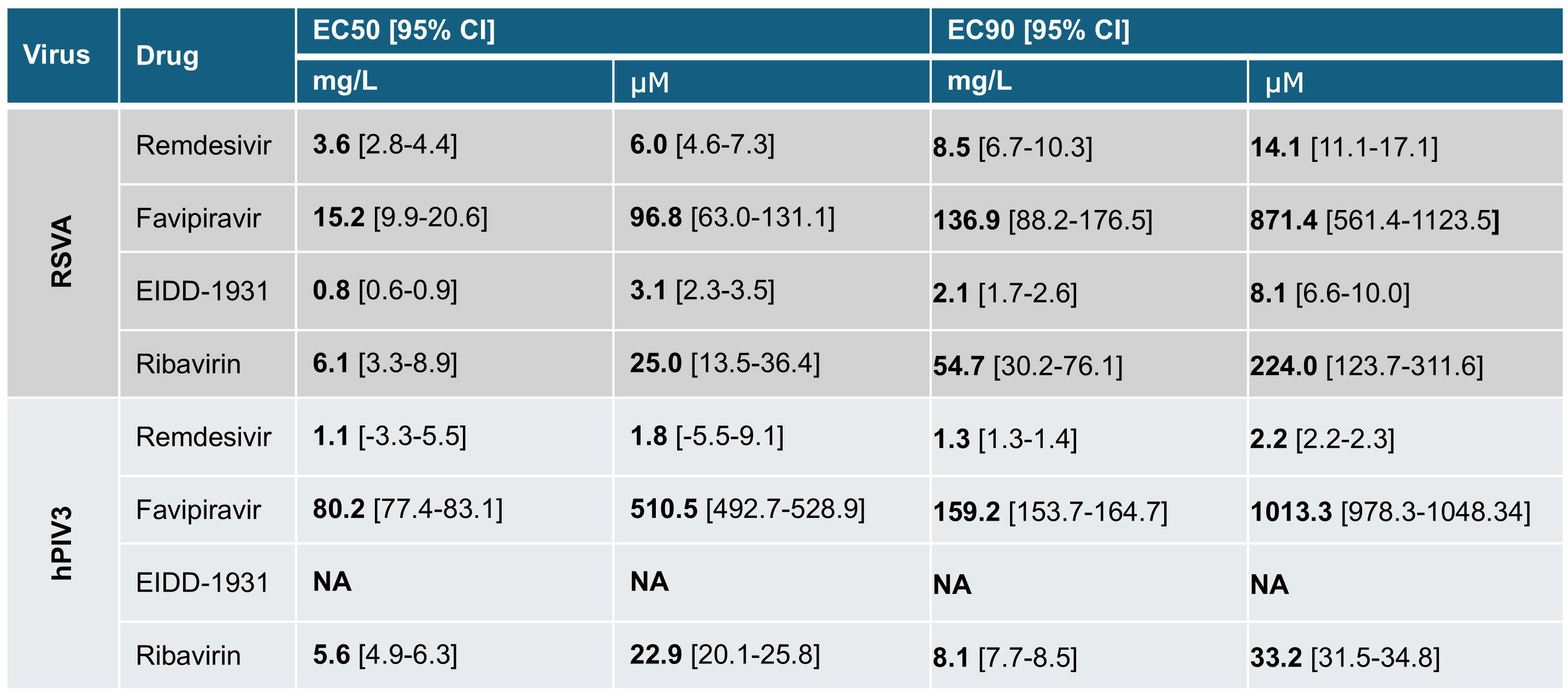
Summary of monotherapy antiviral efficacy.

All four RdRp inhibitors achieved 100% viral inhibition against RSVA within the concentration range tested **[Figure 1E-H]**. For hPIV3, remdesivir, favipiravir and ribavirin all demonstrated effective *in vitro* viral inhibition **[Figure 1I-L]**. Remdesivir demonstrated the overall highest potency against both viruses, with EC50 observed at 1.1mg/L (1.8 μM) and 3.6 mg/L (6.0 μM) against hPIV3 and RSVA, respectively. Additionally, no cytotoxicity was observed throughout the clinically relevant concentrations of remdesivir tested on either cell-line **[Figure 1E&I]**. Ribavirin also demonstrated consistent viral inhibition against RSVA and hPIV3 with EC50s of 6.1 mg/L (25.0 μM) and 5.6 mg/L (22.9 μM), respectively. Ribavirin cytotoxicity was only detected at higher doses, from 128 mg/L (524 μM) for VeroE6 and 32 mg/L (131 μM) for LLC-MK2 **[Figure 1H&L]**.

Favipiravir required substantially higher concentrations to achieve antiviral effects, with EC50s of 15.2 mg/L (96.8 μM) and 80.2 mg/L (510 μM) for RSVA and hPIV3, respectively **[Figure 1F&J]**. For both viruses, significant cytotoxicity was observed at or below the predicted EC90 doses of favipiravir **[Figure 1 & Table 1]**. EIDD-1931 demonstrated the lowest EC50 against RSVA, 0.8 mg/L (3.1 μM) **[Figure 1G]**, but was ineffective against hPIV3 at the concentrations tested. In LLK-MK2 cells, cytotoxicity was evident above 2 mg/L (7.7 μM) with no inhibition of hPIV3 infection **[Figure 1K]**.

### Synergistic dual-drug combinations of remdesivir, favipiravir, molnupiravir and ribavirin against RSVA and hPIV3

We next used our cell-line antiviral assay to evaluate dual-drug combinations of remdesivir, favipiravir, molnupiravir and ribavirin. For each pairwise combination, 4-6 concentrations per drug were selected based on the monotherapy dose-response ranges **[Figures 2&3]**. Concentrations were selected to span the full dynamic range of each single-drug dose response range whilst remaining below the cytotoxicity thresholds in these cells.

**Figure 2.**
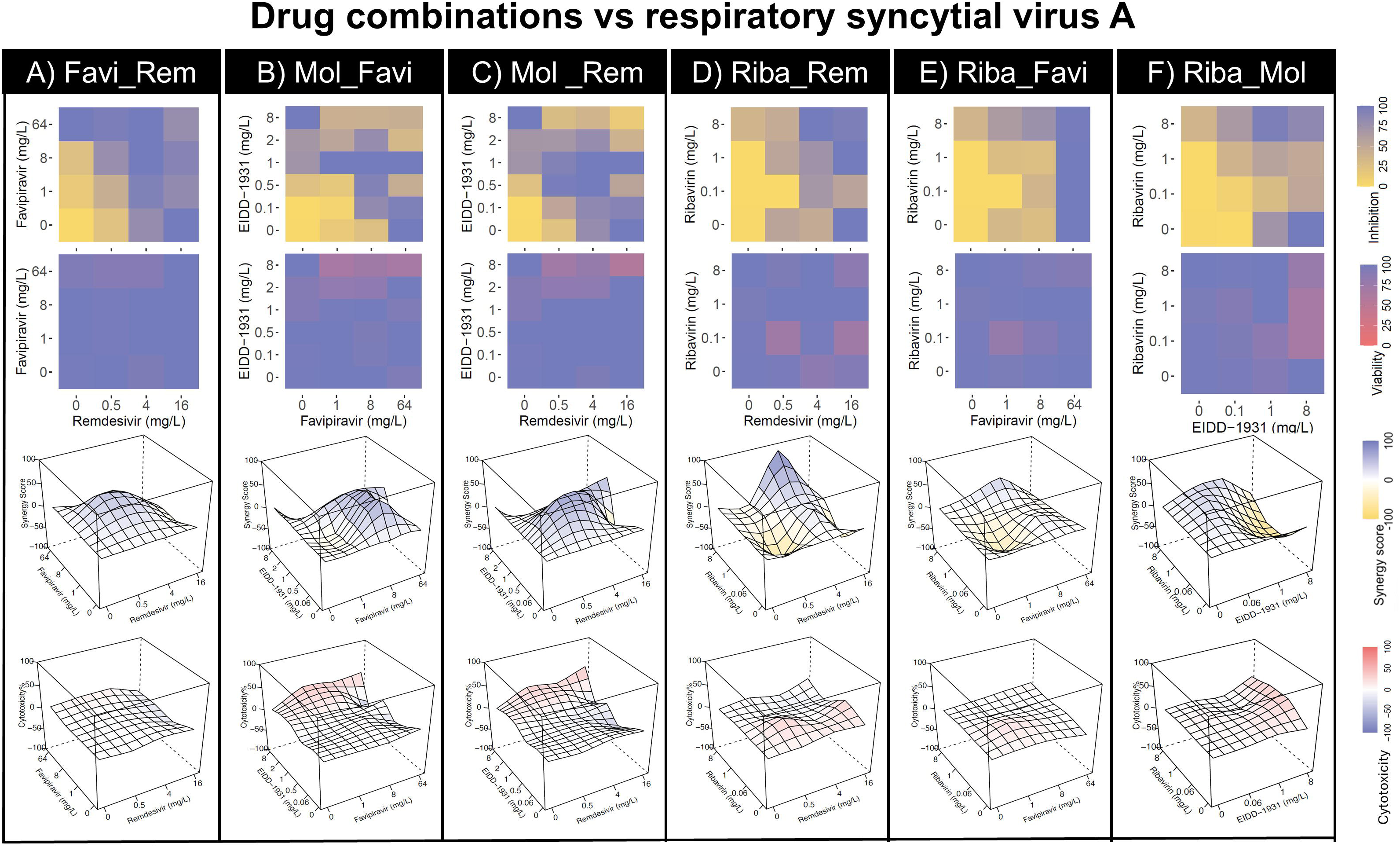
RSVA antiviral combinations and synergy. Dual-drug combinations of antivirals were assayed for viral inhibitory dose response and cytotoxicity for RSVA infection of VeroE6 cells. For each panel (A-F), the dose response grid represents % relative viral inhibition and the cytotoxicity response grid indicates % cell viability. Also shown are 3D contour plots of synergy scores as calculated by the HSA model, alongside cytotoxicity at the corresponding antiviral concentrations. (n=3 per combination) Favi = favipiravir, Rem = remdesivir, Mol = EIDD-1931, Riba = ribavirin.

Remdesivir exhibited synergy with each of the other RdRp inhibitors against RSVA **[Figure 2ACD]** and with all but molnupiravir (which demonstrated no efficacy in monotherapy) against hPIV3 **[Figure 3ACD]**. Synergistic effects were observed at concentrations as low as 0.5 mg/L (0.8 μM) for remdesivir against both viruses. Such combinations were dose sparing, with remdesivir alone having an EC90 of 8.5 mg/L (14.1 μM) against RSVA yet reached a viral inhibition of 97.6% at 0.5 mg/L (0.8 μM) when combined with 0.5 mg/L (1.9 μM) molnupiravir (also below the latter drug’s EC50 of 0.8 mg/L (3.1 μM)). Importantly, these synergistic concentrations were not associated with cytotoxicity.

**Figure 3.**
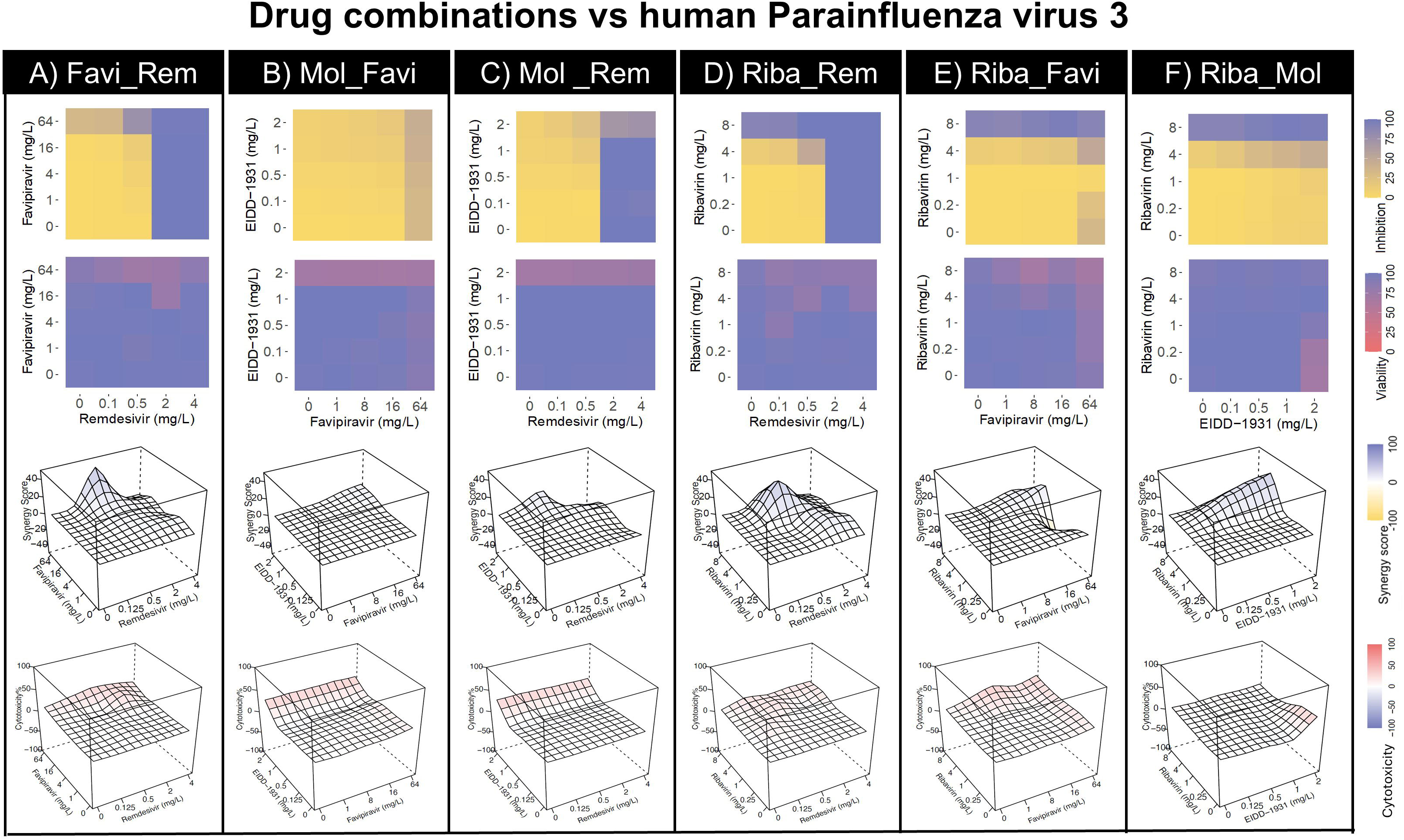
hPIV3 antiviral combinations and synergy. Dual-combinations of antivirals were assayed for viral inhibitory dose response and cytotoxicity for hPIV3 infection of LLC-MK2 cells. For each panel (A-F), the dose response grid represents % relative viral inhibition and the cytotoxicity response grid indicates % cell viability. Also shown are 3D contour plots of synergy scores as calculated by the HSA model, alongside cytotoxicity at the corresponding antiviral concentrations. (n=3 per combination). Favi = favipiravir, Rem = remdesivir, Mol = EIDD-1931, Riba = ribavirin.

Favipiravir also demonstrated synergy at substantially reduced concentrations when combined with remdesivir or molnupiravir against RSVA **[Figure 2AB]**. While its monotherapy EC50 was 15.2 mg/L (96.8 μM), synergy scores reached +62.1 and +60.8 (HSA model) when 8 mg/L (50.9 μM) favipiravir was combined with 0.06 mg/L (0.2 μM) and 0.5 mg/L (1.9 μM) molnupiravir, respectively.

Overall, synergy was less pronounced against hPIV3, with dose responses primarily driven by the higher potency of remdesivir and ribavirin for this virus. The most promising combinations were 0.5 mg/L (0.8 μM) remdesivir with either 64 mg/L (407 μM) favipiravir (HSA score +41.7) or 4 mg/L (16.4 μM) ribavirin (+38.3) **[Figure 3A&D]**.

Across these dual-drug combinations for both RSVA and hPIV3 there was minimal evidence of antagonism. Instances of negative HSA scores were mostly attributable to cytotoxicity at high concentrations, particularly the upper molnupiravir concentrations used in combinations (8 mg/L (30.9 μM) for VeroE6 and 2 mg/L (7.7 μM) for LLC-MK2 cells) **[Figures 2&3]**. A summary of the most synergistic antiviral combinations is provided in **Table 2**.

**Table 2.** Selected combinations of antivirals with greatest synergy in epithelial cell infection assays.

| Virus | Drug A | mg/L | Drug B | mg/L | Viral Inhibition (%) | Synergy Score (HSA model) |
| --- | --- | --- | --- | --- | --- | --- |
| RSVA | Favipiravir | 8 | EIDD-1931 | 0.06 | 82.3 | +62.1 |
| RSVA | Favipiravir | 8 | EIDD-1931 | 0.5 | 92.7 | +60.8 |
| RSVA | Remdesivir | 0.5 | EIDD-1931 | 0.5 | 97.6 | +60.4 |
| RSVA | Remdesivir | 0.5 | EIDD-1931 | 0.06 | 52.0 | +34.1 |
| RSVA | Remdesivir | 0.5 | EIDD-1931 | 1 | 93.0 | +17.9 |
| RSVA | Remdesivir | 0.5 | Favipiravir | 8 | 78.5 | +42.0 |
| RSVA | Remdesivir | 4 | Favipiravir | 1 | 93.7 | +19.3 |
| RSVA | Remdesivir | 0.5 | Favipiravir | 1 | 43.6 | +21.2 |
| hPIV3 | Remdesivir | 0.5 | Favipiravir | 64 | 76.3 | +41.7 |
| hPIV3 | Remdesivir | 0.5 | Ribavirin | 4 | 48.5 | +38.3 |

Blue shading indicates % of monotherapy EC50 concentration Green shading indicated % viral inhibition

### Benchmarking RSVA and hPIV3 infection in a physiologically relevant human airway epithelial model

To benchmark antiviral responses in a more physiologically relevant system, we next performed infection assays using differentiated human airway epithelial cells cultured at air-liquid interface (ALI) **[Figure 4A]**. Cultures were maintained for at least four weeks to achieve full mucociliary differentiation, confirmed by transepithelial electrical resistance (>300 Ω·cm²), and immunofluorescent staining showing intact tight junctions and the presence of mucous-secreting and ciliated epithelial cell types **[Figure 4B-C]**.

**Figure 4.**
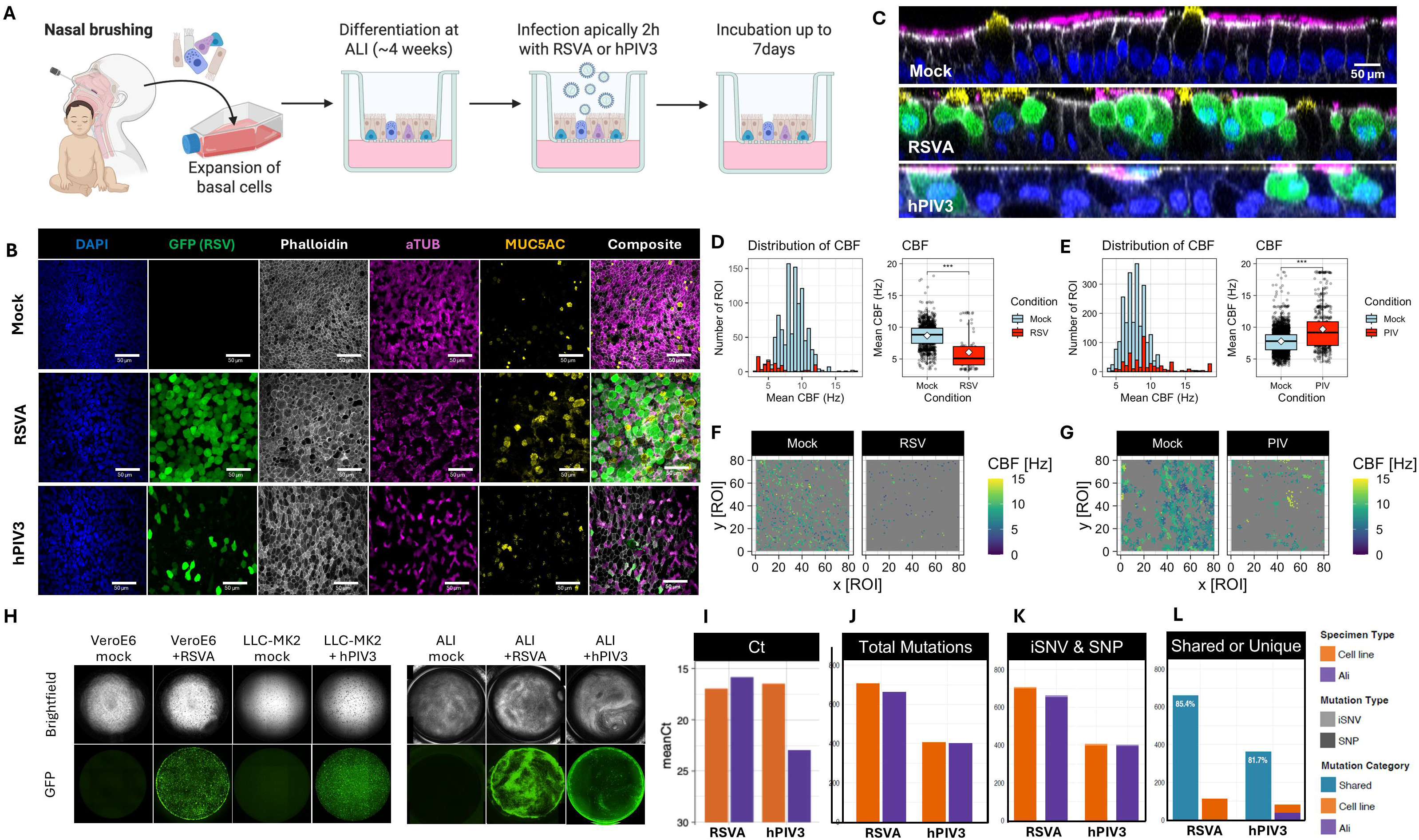
Benchmarking RSVA and hPIV3 infection of differentiated human airway epithelial cell cultures. A. Graphical representation of methodology for culture of primary airway epithelial cell model at air-liquid interface (ALI) and infection with RSVA or hPIV3. B-C. Representative confocal micrographs of primary airway epithelial cells in ALI culture. Cultures were infected with RSVA, hPIV3, or a mock control and fixed for immunofluorescent staining and imaging at 7 days post infection (dpi). Channels represent signal for DAPI staining of cell nuclei (blue), anti-GFP for the GFP-tagged virus (green), phalloidin for tight junctions (white), alpha-tubulin for ciliated epithelial cells (violet), and MUC5AC to indicate mucus-secretory cells (yellow). Orthogonal views (C) are also shown for representative cultures, with the same immunofluorescent staining. D-G. Cilia-beat frequency (CBF) was analysed using high speed video microscopy at 7 dpi with RSVA (D&F) or hPIV3 (E&G) versus mock. Distribution of measured CBF is shown by histogram and box plot (D-E) for each virus. Representative regions of interest (ROI) from video analysis indicate the relative abundance of detectable cilia movement in mocks versus RSVA (F) and hPIV3 (G) infected cultures. H. Representative well scans showing brightfield and GFP signal at 7 dpi for mock and infection conditions with each virus using cell lines (VeroE6 or LLC-MK2 in 96-well format) compared to primary ALI cultures (24-well format). I. The Ct result from qRT-PCR performed on supernatant (for cell lines) or apical wash (for ALI) samples used for viral genomic sequencing, indicating respective viral load at 7 dpi. J-L. Sequencing results for total mutations (J), intrahost single nucleotide variants (iSNV) and single nucleotide polymorphisms (SNP) (K) for each virus from cell line of ALI culture infection model, and the proportion of mutations which were observed in both models or unique (L). Representative images and functional analyses are shown from triplicate ALI cultures per experimental condition.

Both RSVA and hPIV3 established productive infection in the ALI cultures, as determined by GFP fluorescence and microscopy **[Figure 4B-C]**. Despite using the same MOI and time course of infection, RSVA spread across larger, contiguous regions of the epithelium, whereas hPIV3 remained largely confined to individual infected cells **[Figure 4B,C&H]**. This observed difference was quantified by image analysis of GFP-positive well area **[Supplementary Figure 2A]**, and also aligned with relative viral RNA loads measured in apical surface washes by qRT-PCR **[Figure 4I]**. Whole well-scanning fluorescence imaging further illustrated the greater intensity of RSVA replication in ALI cultures compared with mammalian cell lines **[Figure 4H]**. The brightfield scans confirmed preservation of epithelial integrity at 7 dpi, timepoint at which pronounced cytopathology and cell loss are already evident in cell line models **[Figure 4H]**.

High speed video microscopy was used to evaluate ciliary function and the impact of infection on epithelial motility **[Figure 4D-G]**. Mock-infected cultures displayed abundant, rapidly beating cilia, consistent with mature mucociliary differentiation. In contrast, both viruses induced substantial ciliary dysfunction, reflected in altered ciliary beat frequency (CBF) distributions **[Figures 4D&E]**, and representative regions of interest from the video analysis **[Figure 4F&G]**. RSVA infection reduced mean CBF from 8.7 to 5.8 Hz **[Figure 4D&F]**. Consistent with its more limited replication at this timepoint, hPIV3 caused a smaller reduction in the proportion of actively beating cilia. Interestingly, hPIV3 infection also produced localised regions exhibiting abnormally high CBF relative to mock **[Figure 4E&G]**.

Comparative analysis from viral genomic sequencing for cell line and ALI culture samples demonstrated robust concordance of mutagenic patterns **[Figure 4J-L]**. For both RSVA and hPIV3, 85.4% and 81.7% of mutations were shared between the two experimental systems, respectively, at the 7-dpi endpoint. The mutation analysis revealed contributions from both intra-host single nucleotide variants (iSNVs, representing minority variants with allele frequency <50%) and single nucleotide polymorphisms (SNPs, representing consensus-level variants with allele frequency >50%), with the iSNV representing around 1% of the total mutations. The predominance of shared mutations across experimental models provides a robust baseline for the following analysis of the additional drug-induced mutagenic effects.

### RdRp inhibitors reduce viral replication and preserve functional cilia activity during infection of the primary airway epithelial model

Concentrations of the RdRp inhibitor antivirals were selected for testing in the primary airway model based on the dual-drug combinations that’s showed the strongest synergy in cell line assays, along with the corresponding single drug doses. Antivirals were delivered via the basolateral media component of the ALI cultures. Inhibition of viral replication was determined from viral-GFP data **[Supplementary Figure 2B&C]**, used to calculate relative viral inhibition in live cultures and subsequently validated by post-fixation immunofluorescent staining, for both RSVA **[Figure 5]** and hPIV3 **[Figure 6]**.

**Figure 5.**
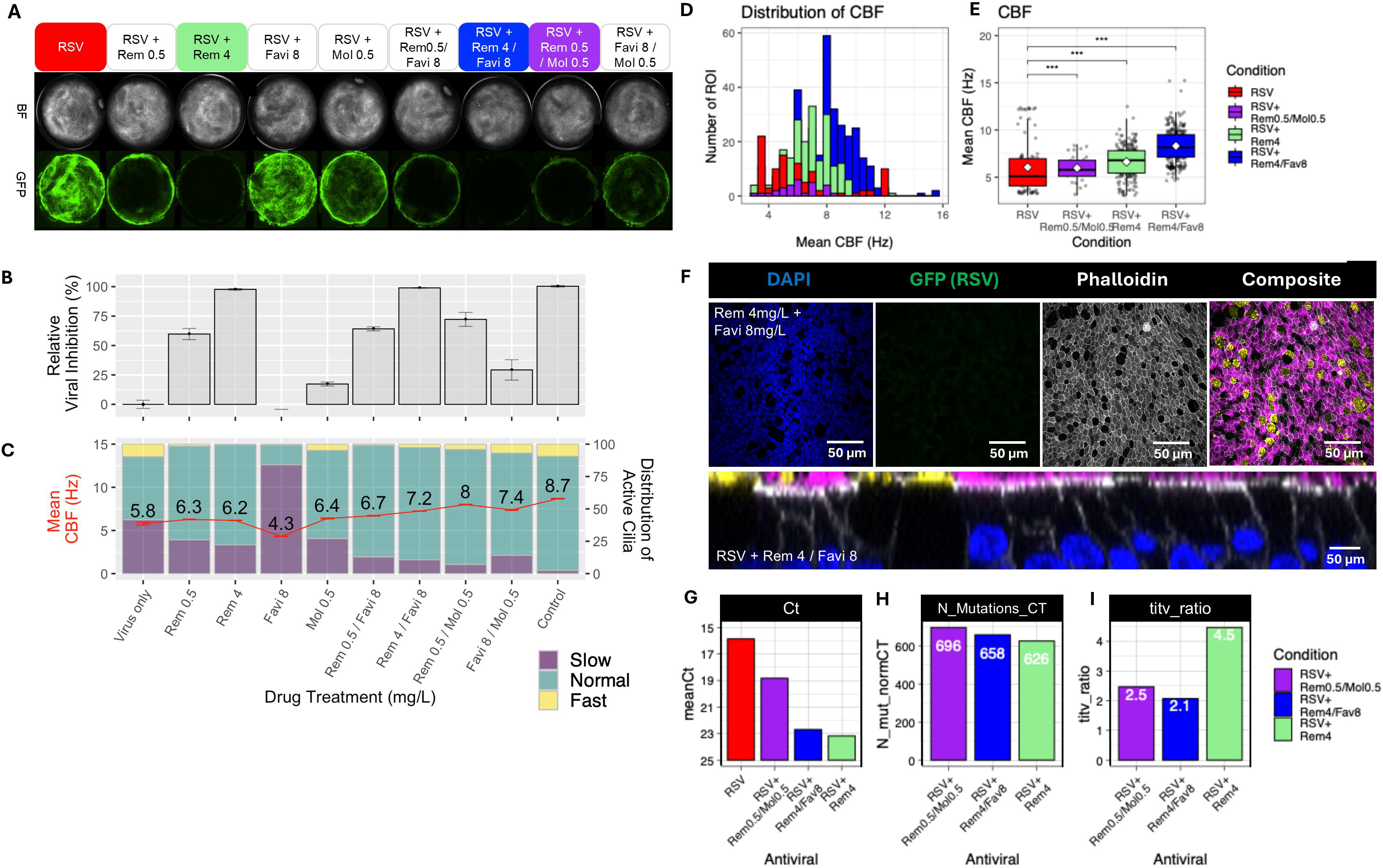
Efficacy of antiviral combinations on RSVA infection of primary airway epithelial cell cultures. A. Representative micrographs of primary airway epithelial cells cultured at air-liquid interface for infection assays with RSVA (A), with whole-well brightfield (BF) and viral GFP fluorescence shown at 7 days post-infection (dpi). B. GFP intensity was quantified by image analysis to calculate relative viral inhibition for all condition. C. High speed video microscopy was used to analyse cilia beat frequency (CBF) for all conditions, with the red line indicating the mean CBF in Hz. D-E. Histogram and box plot representation of CBF distribution for RSVA only and selected antiviral drug conditions. F. Representative confocal micrographs of RSVA-challenged ALI culture with 4 mg/L remdesivir and 8 mg/L favipiravir combination, fixed for immunofluorescent staining and imaging at 7 dpi. Channels represent signal for DAPI staining of cell nuclei (blue), anti-GFP for the GFP-tagged virus (green), phalloidin for tight junctions (white), alpha-tubulin for ciliated epithelial cells (violet), and MUC5AC to indicate mucus-secretory cells (yellow). Orthogonal view also shown for representative culture, with the same immunofluorescent staining. G. Ct result from RSVA qRT-PCR performed on apical wash samples collected for RSVA viral genomic sequencing, indicating respective viral load at 7 dpi. H. Observed mutation rate with selected antiviral conditions, normalised against Ct. I. The Transition/Transversion (titv) ratio of mutation analysis with selected antiviral conditions. Representative images and functional analyses are from triplicate ALI cultures per experimental condition. Favi = favipiravir, Rem = remdesivir, Mol = EIDD-1931

**Figure 6.**
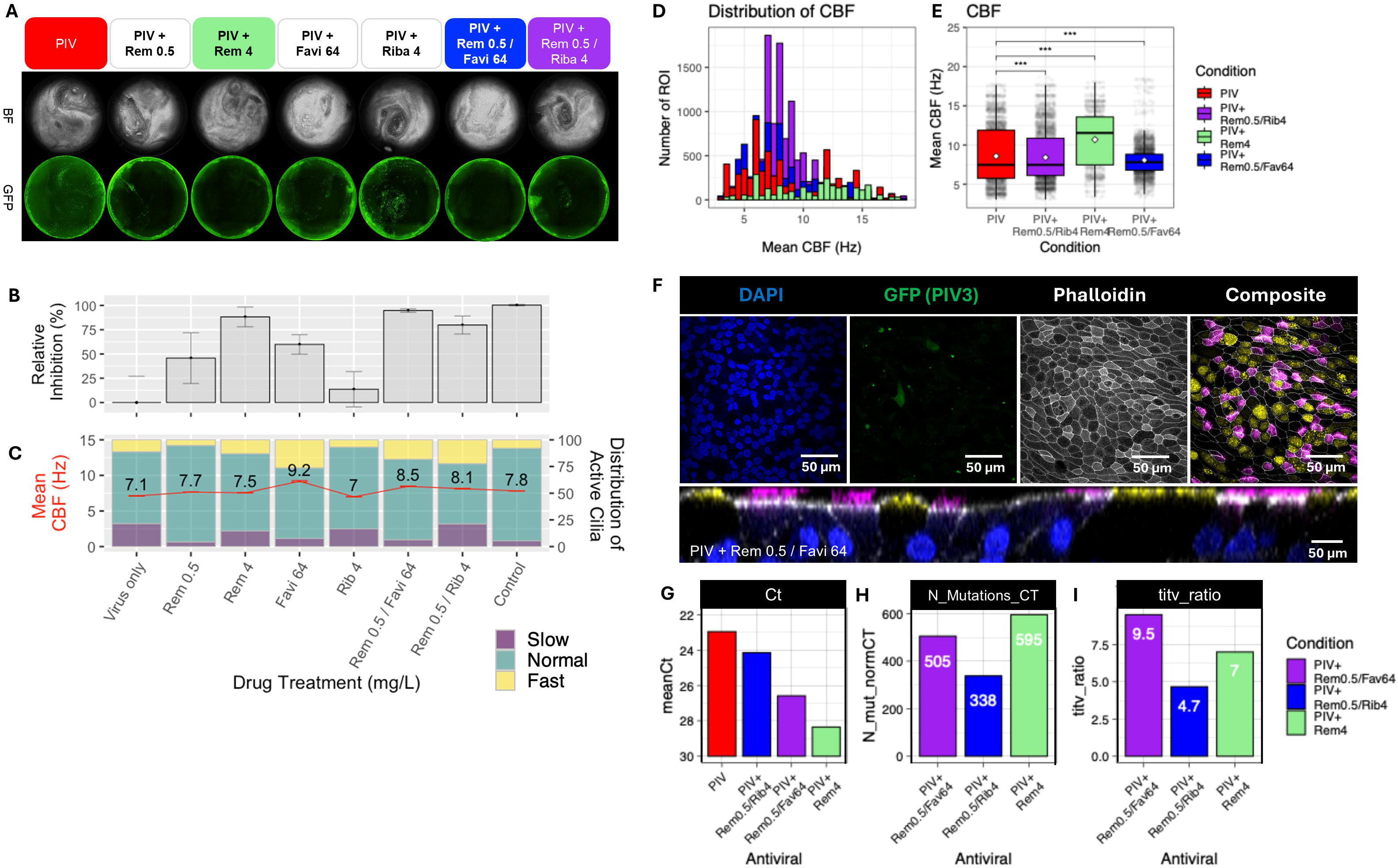
Efficacy of antiviral combinations on hPIV3 infection of primary airway epithelial cell cultures. A. Representative micrographs of primary airway epithelial cells cultured at air-liquid interface for infection assays with hPIV3 (A), with whole-well brightfield (BF) and viral GFP fluorescence shown at 7 days post-infection (dpi). B. GFP intensity was quantified by image analysis to calculate relative viral inhibition for all condition. C. High speed video microscopy was used to analyse cilia beat frequency (CBF) for all conditions, with the red line indicating the mean CBF in Hz. D-E. Histogram and box plot representation of CBF distribution for hPIV3 only and selected antiviral drug conditions. F. Representative confocal micrographs of hPIV3-challenged ALI culture with 0.5 mg/L remdesivir and 64 mg/L favipiravir combination, fixed for immunofluorescent staining and imaging at 7 dpi. Channels represent signal for DAPI staining of cell nuclei (blue), anti-GFP for the GFP-tagged virus (green), phalloidin for tight junctions (white), alpha-tubulin for ciliated epithelial cells (violet), and MUC5AC to indicate mucus-secretory cells (yellow). Orthogonal view also shown for representative culture, with the same immunofluorescent staining. G. Ct result from hPIV3 qRT-PCR performed on apical wash samples collected for hPIV3 viral genomic sequencing, indicating respective viral load at 7 dpi. H. Observed mutation rate with selected antiviral conditions, normalised against Ct. I. The Transition/Transversion (titv) ratio of mutation analysis with selected antiviral conditions. Representative images and functional analyses are from triplicate ALI cultures per experimental condition. Favi = favipiravir, Rem = remdesivir, Riba = ribavirin.

Overall, the RdRp inhibitors displayed broadly similar level of efficacy in the primary airway model compared to the cell line assays. For RSVA, the combination of 0.5 mg/L (0.8 μM) remdesivir with 8 mg/L (50.9 μM) favipiravir achieved 64.2% inhibition in the ALI cultures model compared to 78.5% in VeroE6 cells **[Figures 5A&B]**. Against hPIV3, 0.5 mg/L (0.8 μM) remdesivir in combination with 64 mg/L (407 μM) favipiravir reduced infection by 94.8% in the ALI model versus 76.3% inhibition in LLC-MK2 cells **[Figures 6A&B]**. RSVA infection caused marked epithelial disruption, including loss of epithelial tight junction integrity identified by phalloidin staining, whereas these structural changes were prevented by effective antiviral concentrations such as 4 mg/L (6.6 μM) remdesivir and 8 mg/L (50.9 μM) favipiravir **[Figure 5F]**.

Remdesivir showed enhanced efficacy in the ALI model for both viruses. Whilst the cell line EC50s were 3.6 mg/L (6.0 μM) for RSVA and 1.1 mg/L (1.8 μM) for hPIV3 **[Table 1]**, a substantially lower dose of 0.5 mg/L (0.8 μM) was sufficient for 59.7% and 45.8% viral inhibition in ALI cultures **[Figures 5B & 6B]**. In contrast, 8 mg/L (50.9 μM) favipiravir alone did not inhibit RSVA infection, and did not enhance the efficacy of remdesivir or molnupiravir at the concentrations tested. However, 64 mg/L (407 μM) favipiravir produced a 59.9% reduction in hPIV3 infection (versus an EC50 of 80.2 mg/L (510 μM) in LLC-MK2 cells) and showed modest synergy in combination with 0.5 mg/L (0.8 μM) remdesivir **[Figure 6B]**. For both viruses, antiviral concentrations which caused strong inhibition of replication as determined by viral GFP also resulted in reduced viral load in apical washes, and were associated with accumulation of drug-induced viral genomic mutations **[Figure 5G-I, 6G-I]**.

High speed video microscopy was further used to analyse whether antiviral treatment could prevent virus-induced ciliary disruption **[Figures 5D-E & 6D-E]**. RSVA infection caused both loss of motile cilia and a significant reduction in mean CBF, but this was attenuated by antiviral drug combinations that produced high levels of viral inhibition such as 4 mg/L (6.6 μM) remdesivir with 8 mg/L (50.9 μM) favipiravir **[Figures 5D&E]**. As hPIV3 alone induced less disruption to CBF than RSVA, the effects of antivirals on cilia activity relative to viral inhibition were less obvious. However, combinations of 0.5 mg/L (0.8 μM) remdesivir with 4 mg/L (16.3 μM) ribavirin or 64 mg/L (407 μM) favipiravir both reduced the proportion of abnormally fast beating cilia previously observed during hPIV3 infection **[Figures 6D&E]**.

### RdRp inhibitors induce signature mutations associated with viral inhibition across infection models

Endpoint samples, consisting of culture supernatant for cell lines or apical surface wash for ALI culture, were collected for qRT-PCR, viral genomic sequencing and mutation analysis. For both RSVA and hPIV3 in both the cell line and primary airway culture models, increasing drug concentrations were associated with a reduction in viral load and inhibition of viral replication **[Figure 7A]**. Consistent with prior reports for molnupiravir and favipiravir in SARS-CoV-2 infection, the total number of viral mutations, corrected for viral load and normalised against mutations occurring over the same time-period in untreated cultures **[Figure 7B]**, increased in a dose-dependent manner^36^.

**Figure 7.**
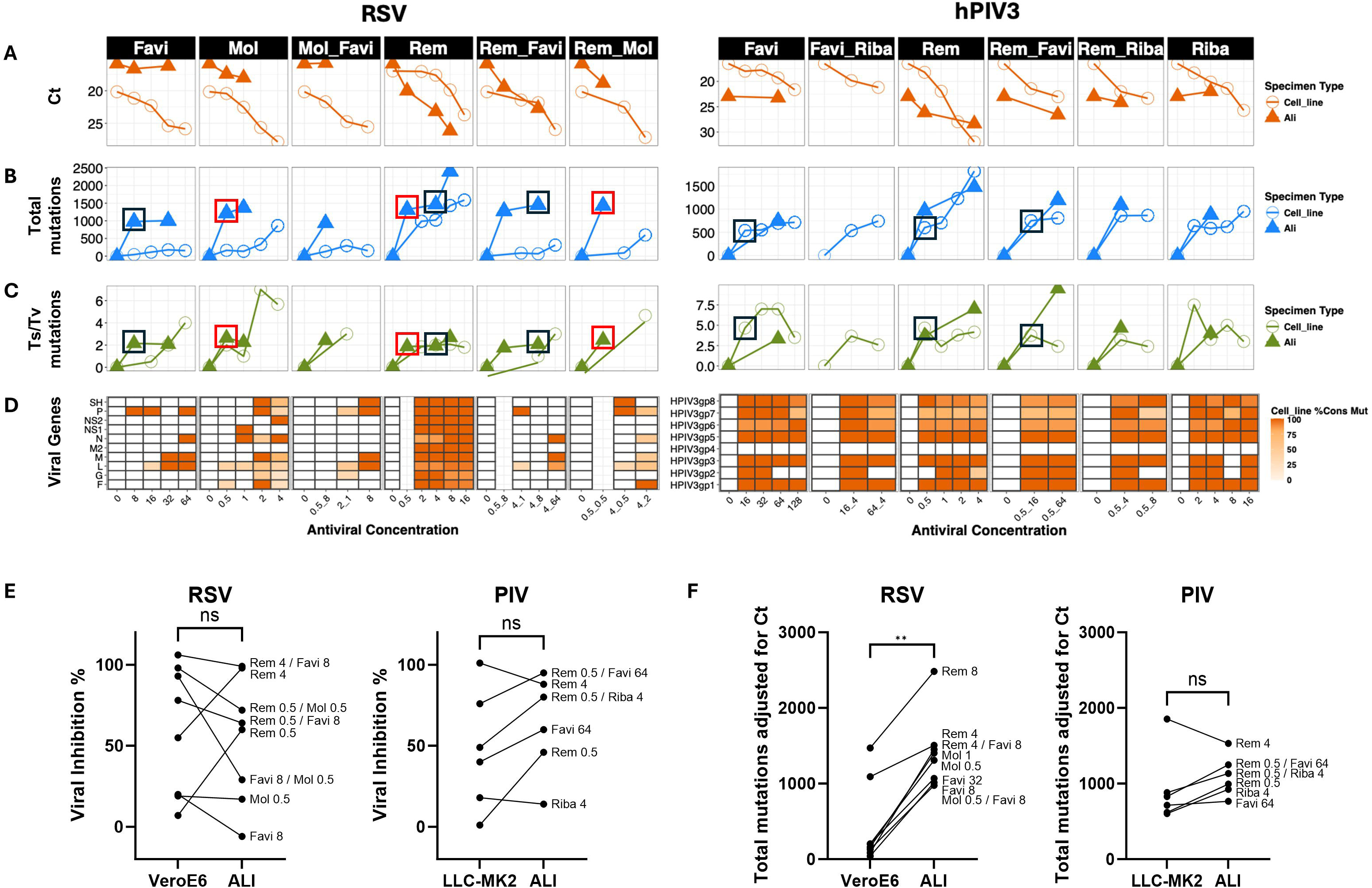
Comparative genomic analysis of antiviral efficacy between cell lines and ALI culture. A. Mean viral load by qRT-PCR CT values across antiviral concentrations for RSVA (left) and hPIV3 (right) in endpoint samples from cell line (orange circles) versus ALI model (orange triangles) infection assays. Lower CT values indicate higher viral replication. B. Total number of mutations compared to zero drug normalized by CT values across antiviral treatments for RSVA (left) and hPIV3 (right) in cell lines (blue circles) and ALI samples (blue triangles). C. Transition/transversion (Ts/Tv) mutation ratios across antiviral treatments for RSVA (left) and hPIV3 (right) comparing cell lines (green circles) and ALI samples (green triangles) values are normalised for CT values and zero drug. Boxes indicate selected corresponding single and dual-combination concentrations for remdesivir-molnupiravir (red boxes) or remdesivir-favipiravir (black boxes). D. Heatmap showing the proportion of consensus mutations across viral genes (y-axis) for different antiviral concentrations (x-axis) in cell line experiments. Colour intensity represents the percentage of consensus mutations (0-100%). E. Comparison of viral inhibition for matching antiviral concentrations between the respective cell line and ALI models for RSVA and hPIV3. F. Comparison of total viral mutations (adjusted for viral load) under matching antiviral concentrations between the respective cell line and ALI models for RSVA and hPIV3.

Unexpectedly, remdesivir was also mutagenic for both RSVA and hPIV3 across the cell lines and ALI models, contrasting with observations in SARS-CoV-2 infection, where remdesivir is generally considered a non-mutagenic chain terminator. As expected for RdRp-targeting nucleoside analogues, transition point mutations (A↔G; C↔T) increased with antiviral treatment, whereas transversion substitutions (purine↔pyrimidine) did not. Accordingly, the transition/transversion (Ts(G-A;C-T) /Tv) ratios increased for all antiviral treatments relative to untreated controls for both RSVA and hPIV3 **[Figure 7C]**.

Notably, the increased viral inhibition seen with dual-drug combination therapies, such as favipiravir (8 mg/L; 50.9 μM) combined with molnupiravir or remdesivir against RSVA, or favipiravir (16 mg/L; 101 μM) combined with ribavirin or remdesivir against hPIV3, was not associated with further increased mutagenesis **[boxed points, Figure 7A-C]**. Across both viruses, mutations were dispersed randomly across the genome, with no clustering, enrichment, or evidence of selection or concentration at known drug-binding or active sites **[Figure 7D]**.

Comparing the two infection systems, the overall level of antiviral inhibition was not significantly different between cell line and ALI models for RSVA (p=0.70) or hPIV3 (p=0.12) **[Figure 7E]**. However, for RSVA, antiviral-induced viral mutagenesis was significantly higher in ALI cultures compared to VeroE6 cells (p=0.0078) **[Figure 7F]**. While hPIV3 showed a minor trend towards higher mutation rates in ALI cultures for most tested antiviral doses, this did not reach statistical significance overall (p=0.22) **[Figure 7F]**.

## Discussion

In this study, we demonstrate that four clinically relevant RdRp inhibitors—remdesivir, ribavirin, favipiravir and molnupiravir—exhibit distinct but complementary antiviral activities against RSVA and hPIV3 across both traditional epithelial cell lines and a physiologically relevant primary airway epithelial ALI model. We show that remdesivir is the most potent monotherapy across viruses and models, that multiple dual-drug combinations act synergistically, and that effective antiviral exposure is associated with dose-dependent increases in transition mutagenesis. Importantly, we further show that antiviral effects and mutagenesis profiles translate into the ALI system, where structural and functional preservation of the epithelium—including maintenance of ciliary beating—provides additional biological validation of therapeutic benefit.^15,37^.

We first characterised the monotherapy activity of each RdRp inhibitor using optimised epithelial cell-line infection assays. The selection of cell lines was driven empirically by the requirement for robust infection and quantifiable CPE or GFP readouts **[Supplementary Figure 1]**. LLC-MK2 cells, a monkey kidney epithelial cell line, were the most permissive for hPIV3 infection, consistent with their extensive use in PIV studies^38^,while VeroE6 cells provided reliable RSVA infection and are well-established in RSV antiviral research^39,40^

Across both viruses, remdesivir emerged as the most effective antiviral tested. We calculated EC50 values for hPIV3 (1.1 mg/L; 1.8 μM) and RSVA (3.6 mg/L; 6.0 μM), which are consistent with or slightly higher than previous reported values using similar strains and assay formats, noting that earlier infection timepoints (72 hours post-infection) used in other studies may yield lower EC50 estimates due to reduced cytopathic contribution^17^.^17^. The efficacy of remdesivir against RSV in alternative cell lines (HEp-2, A549) with EC50 <1 mg/L (<1.7 μM) has also been documented, supporting its broad antiviral potential ^41,42^.

Ribavirin also demonstrated strong antiviral activity against both viruses. Our EC50 for hPIV3 (5.6 mg/L; 22.9 μM) aligns with EC50 values of 2–13 mg/L (8.2-53.2 μM) previously published for both laboratory-adapted and clinical hPIV3 strains^16,17^. Against RSVA, we observed an EC50 of 6.1 mg/L (25.0 μM), comparable to values of 3.9 mg/L (16.0 μM) and 2.7 mg/L (11.1 μM) in HeLa/M and HEp-2 cells^43,44^.

Favipiravir inhibited both RSVA and hPIV3 but required higher concentrations, and cytotoxicity emerged near the EC90 in both VeroE6 and LLC-MK2 cells. Nonetheless, an EC50 of 15.2 mg/L (96.8 μM) for RSVA is within a 10.7-41.0 mg/L (68.1-261 μM) range published in studies using HEp-2 infection models^45,46^. *In vitro* favipiravir efficacy against hPIV infection has been reported at 10.8 mg/L (68.7 μM) and 21.7 mg/L (138 μM) elsewhere, although in different host cells such as Vero-118 and PLC/PRF/5, whilst our EC50 value of 80.2 mg/L (510 μM) in LLC-MK2 cells is likely beyond the clinically achievable range ^16,45^.

Molnupiravir (using the active metabolite EIDD-1931) produced the strongest antiviral effect against RSVA, with an EC50 of 0.8 mg/L (3.1 μM), which is consistent to results between 0.1-1.0 mg/L (0.4-3.9 μM) obtained with clinical RSV strains in HEp-2 cells^47^. In contrast, we observed no inhibition of hPIV3 due to pronounced cytotoxicity above 2 mg/L (7.7 μM) in LLC-MK2 cells, which would prevent any accurate determination of potential hPIV3 inhibition above this concentration. To our knowledge, hPIV sensitivity to molnupiravir has not been reported previously. It is possible that sensitivity of hPIV3 to molnupiravir may be measurable in ALI cultures, if less sensitive to cytotoxicity than LLC-MK2 cells, but that was not tested in this study.

We next evaluated dual-drug combinations, which can provide several advantages over monotherapy including additive or ideally synergistic inhibition of viral replication^21^. Recently the use of combined antivirals has been investigated in greater detail in the context of SARS-CoV-2 therapy, including demonstrated synergism in cell culture and animal models^9,24,25^. In a directly relevant clinical setting, a case series at our partner hospital, Great Ormond Street Hospital for Children, has been recently published outlining potential benefits from combination favipiravir, ribavirin, and nitazoxanide to treat respiratory viral infections in severe T-cell deficient children^48^.

Consistent with findings in SARS-CoV-2 models, we observed strong synergy for several RdRp inhibitor pairs against RSVA *in vitro*— favipiravir-molnupiravir, remdesivir-molnupiravir, and remdesivir-favipiravir. Against hPIV3 infection, synergy was more limited but evident for remdesivir–favipiravir and remdesivir-ribavirin. These combinations were also dose-sparing: for example, remdesivir alone required 8.5 mg/L (14.1 μM) to achieve an EC90 against RSVA, yet 0.5 mg/L (0.8 μM) in combination with 0.5 mg/L (1.9 μM) molnupiravir achieved 97.6% inhibition. Importantly, synergistic doses showed minimal cytotoxicity. Synergistic combinations of RdRp inhibitors have previously been shown for SARS-CoV-2, including using the *in vivo* Syrian hamster model ^25,49,50^. Radoshitzky et al. have also shown a broad-spectrum potential of remdesivir against many viral classes with no antagonism with a favipiravir combination^51^.

Our mutation analysis parallels earlier work showing that transition mutagenesis (A↔G; C↔T) increases with nucleoside analogue exposure and can serve as a biomarker for antiviral activity against norovirus, influenza B, SARS-CoV-2 and RSVA^36,52-54^. As expected, favipiravir and molnupiravir induced increasing transition mutations after correcting for viral load and normalising to untreated controls. Unexpectedly, remdesivir, which is not reported as directly mutagenic as with favipiravir or molnupiravir in SARS-CoV-2, also induced significant increases in total and transition mutations in both RSVA and hPIV3 in both infection models^36^. Our data, which are the first to examine potential mutagenic signatures in this way for RSVA and hPIV3, raise the possibility of previously unrecognised virus-specific properties of remdesivir’s mechanism of action in paramyxoviruses.

It has been previously observed in SARS-CoV-2 infected hamsters that combination therapy modestly increased total mutation burden compared with each drug alone^36^. While the current study was not designed to measure that effect, we did observe a modest increase in total mutation burden for favipiravir plus remdesivir for hPIV3 in LLC-MK2 cells, although this was less obvious in RSVA infected ALI cultures **[Figure 7B&C]**. A targeted study would be required to determine whether mutagenesis contributes mechanistically to synergy.

The primary airway ALI system provided several advantages over standard monolayer cell lines in this study. First, unlike VeroE6 and LLC-MK2, which supported RSV or hPIV3 but not both, ALI cultures supported infection by both viruses, allowing direct cross-virus comparison under identical culture conditions^27^. Second, the mammalian cell lines have limited biological relevance to human infection, and do not display important respiratory epithelial characteristics such as transport proteins, mucus secretion and active motile cilia^28^.

Studies on respiratory viruses have begun to utilise these advantages for diverse research aims, such as investigating the impact of tissue donor age on RSV infection kinetics, the mutation accumulation rates when passaging hPIV, and the cell tropism and epithelial responses to SARS-CoV-2 infection ^31,55,56^. Respiratory ALI models are also emerging as a viable tool for improved antiviral testing. ALI culture of human nasal epithelial cells has been applied for screening antiviral nucleosides for SARS-CoV-2 inhibition^25,57^. We have also previously demonstrated the potential application of a 96-HTS Transwell ALI model for high-throughput screening with RSV^58^.

Overall, the effective doses determined from cell line assays for the RdRp inhibitors used in our study translated into comparable viral inhibition in the ALI infection model, although enhanced efficacy of remdesivir and reduced efficacy of favipiravir were observed for RSVA versus the VeroE6 results. Limited published studies have compared antiviral efficacy between such models. Mirabelli et al. previously determined EC50s for ribavirin and other replication or fusion inhibitors against RSV using HEp-2 cells, and then demonstrated their efficacy with fully differentiated human airway epithelium cells at ALI, although this study differed in delivering higher doses (10-100x EC50) at later timepoints (1-5 dpi) to mimic therapeutic regimens^44^. Our data further aligns with the findings of Lin et al. for hPIV infection models. They also reported synergy for combinations of remdesivir and ribavirin in LLC-MK2 cells, as well as investigating ribavirin alone and in combination with GS-441524, the parent nucleoside of remdesivir, in infections of human nasal airway epithelial ALI cultures^17^. While viral inhibition was comparable between the models, we did observe higher drug-associated mutation rates for RSVA using primary airway cultures. While there is some evidence that ALI cultures are associated with lower viral mutation rate during routine passaging^56^, it is also established that these primary airway cells reproduce complex innate immunity and antimicrobial responses and secretions which could influence the mutation pressure during viral replication^31,59^. The specific mechanisms behind an increased antiviral drug-induced mutagenesis for RSVA in this model requires further study. Yet this finding does suggest these antivirals have greater potential mutagenic activity in the more physiologically-relevant differentiated airway model, with implications for their *in vivo* applications.

### Limitations

This study acknowledges several limitations. Firstly, comparing antiviral activity across different cell systems—including immortalized lines and ALI cultures—introduces variability driven by differences in viral entry receptors, innate signalling, and cellular responses, all of which can influence drug potency. Cytotoxicity further constrained concentration ranges for some antivirals (most notably for molnupiravir in LLC-MK2 cells), limiting full dose–response characterization for hPIV3. These factors highlight the need for physiologically relevant models such as ALI cultures, while also emphasizing the importance of broader standardization across laboratories to improve reproducibility

Secondly, *in vitro* findings may not fully predict *in vivo* therapeutic efficacy, pharmacokinetics, or host immune responses. Future studies using small animal models or clinical data will be essential to confirm the translational relevance of the synergistic combinations identified here.

Finally, while mutagenesis analysis provided important mechanistic insights, sequencing was performed at single endpoint timepoints. Time-resolved analysis would better characterise mutation accumulation, distinguish lethal mutagenesis from other inhibitory processes, and clarify whether mutagenic synergy contributes directly to antiviral synergy.

## Conclusions

Our results show that combinations of RdRp inhibitors suppress RSVA and hPIV3 replication, with remdesivir displaying the greatest potency and several combinations exhibiting strong synergy, especially remdesivir-favipiravir, remdesivir-molnupiravir, and favipiravir-molnupiravir. These effects translated into primary airway epithelial cultures, where effective antiviral concentrations preserved epithelial integrity and ciliary function after viral challenge. RdRp inhibitor–associated mutagenesis increased dose-dependently across models, including unexpected mutagenic signatures for remdesivir in RSVA and hPIV3, revealing potential virus-specific mechanisms. These findings strengthen the rationale for combination RdRp therapy for respiratory RNA infections, particularly in vulnerable patient groups, and as a broad-spectrum option for future viral threat preparedness.

## Acknowledgements

This work was funded by Great Ormond Street Children’s Charity (V4022), The John Black Charitable Foundation, and the NIHR Great Ormond Street Hospital Biomedical Research Centre.

C.M.S. acknowledges support from a UKRI/BBSRC research grant (BB/V006738/1) and Animal Free Research UK (AFR19-20274), who supported M.P. JB receives funding from the NIHR UCL/UCLH Biomedical Research Centre. The funders had no role in study design, data collection and analysis, decision to publish or preparation of the manuscript. The views expressed are those of the author(s) and not necessarily those of the NHS, the NIHR or the Department of Health.

We acknowledge and thank UCLGenomics (RRID:SCR_027010) for undertaking viral whole genome sequencing. We also thank the Cell Services science technology platform (STP) at the Francis Crick Institute, London, UK, for providing the African green monkey kidney cell line Vero E6 (ATCC: CVCL_0574 authenticated for use in this study).

## Author Contributions

SE, AIJ and JB designed the study. SE and AIJ, conducted experiments, analysed data, and cowrote the manuscript. CMS, SZ, JB and LB performed data analysis and contributed to manuscript and figures. MW performed microscopy experiments and analysis. JA, MP, EKH, HM, and MT supported viral and cell culture and *in vitro* experiments. HT and RW carried out and oversaw viral genome sequencing. CM and AK supported qRT-PCR experiments and sample processing. COC provided support through ethics and donor recruitment for primary airway cultures. JB conceived the study and obtained funding. JB, CMS, and JFS oversaw data analysis and interpretation, and the write-up of the manuscript.

## Figure Legends

**Supplementary Figure 1. Determining optimal epithelial cell line for RSVA and hPIV3 infection assays.** A-B. Three epithelial cell lines VeroE6, Calu3 and LLC-MK2 were screened for the development of quantifiable infection markers with RSVA (A) and hPIV3 (B). For each pairing of virus and cell type, infections were performed at MOIs of 0.1, 0.02 or 0.004. Infection progression was determined by either cytopathic effects (CPE) using quantification of endpoint crystal violet staining of surviving cells at each timepoint, or from fluorescence intensity measurements of viral-GFP signal. Measurements were made at timepoints of 3-, 5- or 7-days post-infection. C-D. The extent of CPE and viral-GFP was compared for each cell and virus pairing at a 7 days post-infection endpoint at either an MOI of 0.004 (C) or 0.1 (D). n=3.

**Supplementary Figure 2. Viral GFP quantification for primary airway culture infections.** Fluorescence micrograph image analysis was used to calculate the percentage area of each well with positive GFP fluorescence, as a quantification of viral infection. A. Comparison of virus only condition for RSVA and hPIV3. B-C. Quantification of viral GFP with and without antiviral drugs for RSVA (B) and hPIV3 (C). Asterisks indicate conditions with a significantly reduction versus virus only control. C. All values shown are mean (±SEM) of triplicate cultures. * p<0.05, ** p<0.01, *** p<0.001, **** p<0.0001.

## Notes

### Competing Interest Statement

The authors have declared no competing interest.

