## Supplementary figures and images for "Antiviral drug synergy and mutational signatures in different epithelial cell models of RSV and hPIV infection"

### Supplementary Figure 1

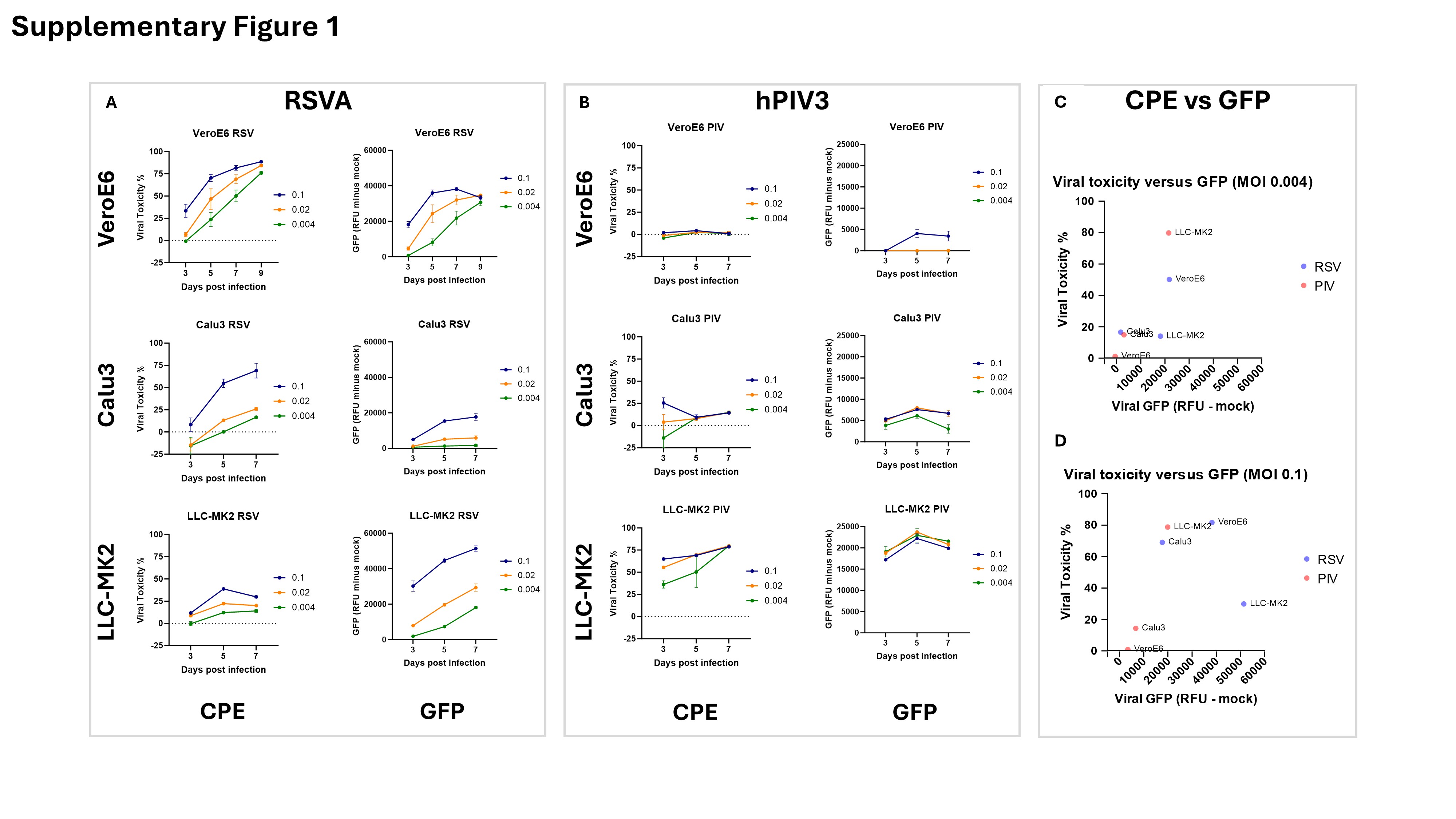

### Supplementary Figure 2

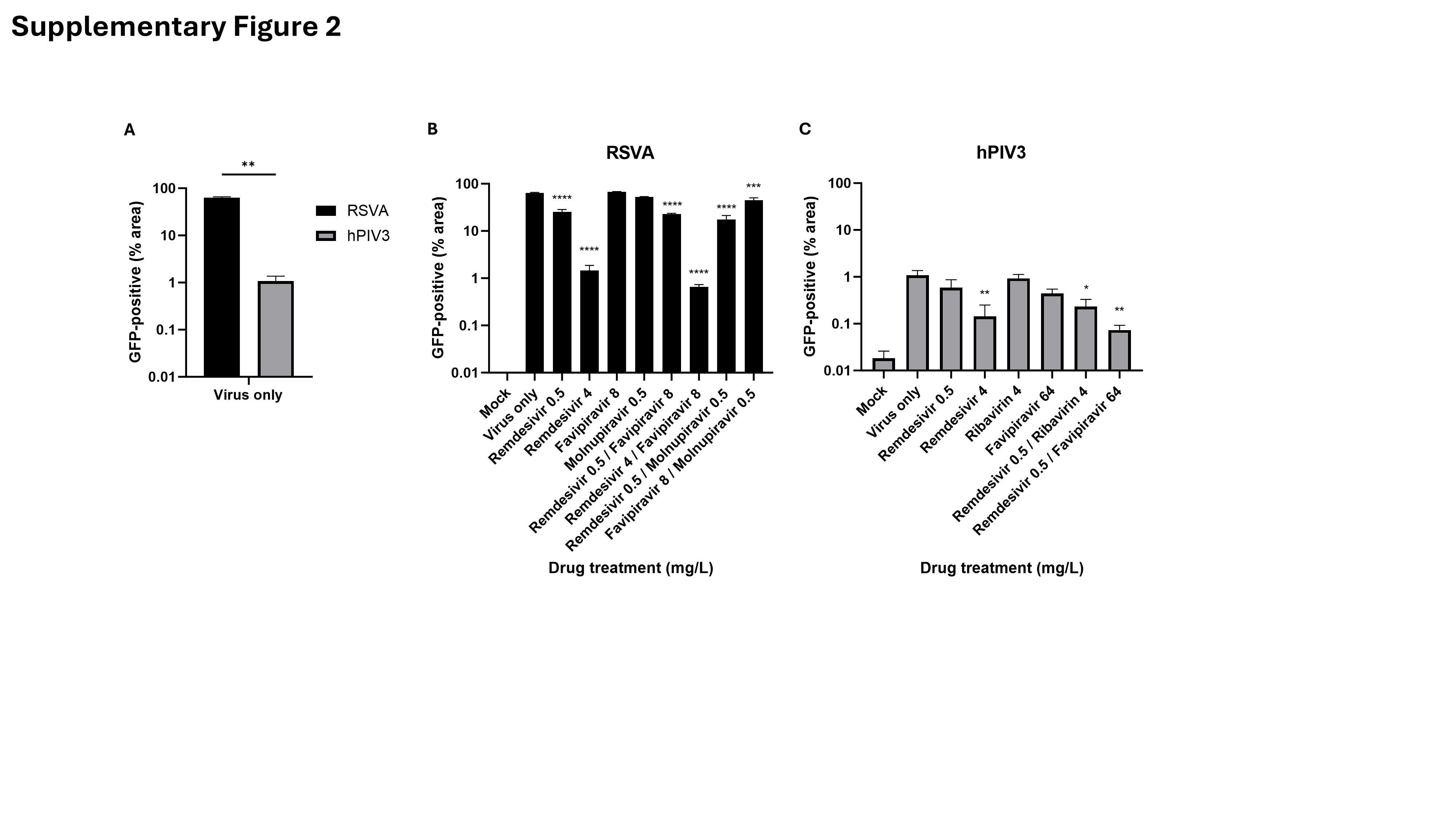
